# Biochemical characterization and essentiality of *Plasmodium* fumarate hydratase

**DOI:** 10.1101/158956

**Authors:** Vijay Jayaraman, Arpitha Suryavanshi, Pavithra Kalale, Jyothirmai Kunala, Hemalatha Balaram

## Abstract

*Plasmodium falciparum* (Pf), the causative agent of malaria has an iron-sulfur cluster-containing class I fumarate hydratase (FH) that catalyzes the interconversion of fumarate to malate, a well-known reaction in the tricarboxylic acid cycle. In humans, the same reaction is catalyzed by class II FH that has no sequence or structural homology with the class I enzyme. Fumarate, generated in large quantities in the parasite as a byproduct of AMP synthesis is converted to malate by the action of FH, and subsequently used in the generation of the key metabolites oxaloacetate, aspartate and pyruvate. Here we report on the kinetic characterization of purified recombinant PfFH, functional complementation of *fh* deficiency in *Escherichia coli* and mitochondrial localization in the parasite. The substrate analog, mercaptosuccinic acid was found to be a potent inhibitor of PfFH with a K_i_ value in the nanomolar range. Knockout of the *fh* gene was not possible in *P. berghei* when drug-selection of the transfectants was performed in BALB/c mice while the gene was amenable to knockout when C57BL/6 mice were used as host, thereby indicating mouse-strain dependent essentiality of the *fh* gene to the parasite.

---

*Plasmodium falciparum* (Pf), the causative agent of the most lethal form of malaria, during its intraerythrocytic asexual stages, derives ATP primarily from glycolysis with low contribution from mitochondrial pathways (1, 2). The bulk of pyruvate formed is converted to lactic acid with a minor amount entering the tricarboxylic acid (TCA) cycle, the flux through which is upregulated in sexual stages (2). Key intermediates that anaplerotically feed into the TCA cycle are α-ketoglutarate derived from glutamate, oxaloacetate (OAA) from phosphoenolpyruvate, and fumarate from adenosine 5’-monophosphate (AMP) synthesis. Synthesis of AMP in the parasite is solely from inosine-5’-monophosphate (IMP) through a pathway involving the enzymes adenylosuccinate synthetase (ADSS) and adenylosuccinate lyase (ASL). The net reaction of ADSS and ASL involves consumption of GTP and aspartate and, generation of GDP, P_i_ and fumarate. In the rapidly dividing parasite with an AT-rich genome and high energy requirements, leading to a high demand for adenine pools, one would expect a high flux of fumarate generation. The parasite does not secrete fumarate but instead, the carbon derived from this metabolite can be traced in malate, OAA, aspartate, pyruvate (through PEP) and lactate (3). The metabolic significance of this fumarate anaplerosis is still obscure. In this context, fumarate hydratase (FH, fumarase) the key enzyme to metabolize fumarate becomes an important candidate for further investigation.

Fumarate hydratase (fumarase, E.C. 4.2.1.2) catalyzes the reversible conversion of fumarate to malate. The stereospecific reaction involves the *anti-addition* of a water molecule across the carbon-carbon double bond of fumarate resulting in the formation of S-malate (L-malate). The reverse reaction proceeds with the elimination of a molecule of water from malate in an anti-fashion (4–6). FH comes in two biochemically distinct forms; class I FH a thermolabile, oxygen sensitive, 4Fe-4S cluster containing enzyme and, class II FH, a stable, oxygen insensitive and iron-independent enzyme (7). Class I FH is further divided into two types, two-subunit and single-subunit, depending on the number of genes that encode the functional enzyme (8). There is no sequence homology between these two classes of enzymes. Class I fumarases display substrate promiscuity; apart from catalyzing the interconversion of fumarate and malate, these enzymes also interconvert S, S-tartrate and oxaloacetate and, mesaconate and S-citramalate with varying catalytic efficiencies (9, 10). The 4Fe-4S cluster is bound to the enzyme by 3 metal-thiolate bonds formed between 3 conserved cysteine residues in the protein and 3 ferrous ions (11). The fourth iron in the cluster proposed to be held loosely by a hydroxyl ion is thought to be directly involved in substrate binding and catalysis as seen in the enzyme aconitase(12, 13).

Both classes of FHs are distributed in all three domains of life with class I FH being more prevalent in archaea, prokaryotes and lower eukaryotes. Many organisms have genes corresponding to both the classes, as in *Escherichia coli* (Ec), which has three FH encoding genes *viz., fum A, B and C.* Fum A and fum B are 4Fe-4S cluster containing class I enzymes, while Fum C belongs to class II type FH. Recently, another gene *fum D* has been identified in the *E. coli* genome to code for a class I fumarase with altered substrate preferences (14).

The structural and biochemical characteristics of class II FH are thoroughly studied from different organisms *viz.,* human, porcine, yeast, *E. coli* and other sources (15–19). On the other hand, class I FH is not well studied owing to its thermolabile and oxygen sensitive nature. All Apicomplexans and Kinetoplastids possess only class I FH, whereas Dinoflagellates have both the classes (20). Biochemical characterization of class I FH from *Leishmania major* (Lm) and *Trypanosoma cruzi,* both Kinetoplastids (21, 22), and the 3-dimensional structure of LmFH II (11) are the only reports of class I FH from eukaryotes. All *Plasmodium* species have one gene annotated putatively as fumarate hydratase that remains to be characterized. Genetic investigations on the role of TCA cycle enzymes in *P. falciparum* have revealed non-essentiality of all genes of TCA cycle except FH and malate-quinone oxidoreductase (23). Recently, a metabolic network reconstruction of pathways in artemisinin resistant *P. falciparum* strains has identified FH reaction as uniquely essential to these parasites (24). Biochemical characterization of PfFH could throw light on unique features of the enzyme and also provide leads for the development of inhibitors.

We report here the kinetic characterization and substrate promiscuity of PfFH, studied using *in vitro* assays on the recombinant enzyme and *E. coli* based functional complementation. DL-mercaptosuccinic acid (DL-MSA), a malate analog was found to be a competitive inhibitor of the *P. falciparum* enzyme. DL-MSA inhibited the growth of the Δ*fumACB strain* of *E. coli* expressing PfFH as well as the asexual intraerythrocytic stages of *P. falciparum* in *in vitro* cultures. Attempts at generating *fh* null *P. berghei* grown in BALB/c mice yielded drug resistant clonal populations that had retained the *fh* gene, implying its essentiality. However, *fh* gene knockout was obtained when the parasites were grown in C57BL/6 mice. This suggests mouse-strain dependent essentiality of the *fh* gene in *P. berghei.*

## RESULTS AND DISCUSSION

### Distribution of Class I fumarate hydratase in eukaryotes

Although both class I and class II FHs catalyze the conversion of fumarate to malate, it is the class II FHs that are widely distributed across eukaryotic organisms. To elicit possible correlations between the presence of class I FH and the nature of the organisms such as their uni-or multi-cellularity and parasitic or free-living lifestyle, eukaryotes with class I FH were catalogued (Table 1). In addition, the presence/absence of 1) class II FH in organisms having class I FH and 2) mitochondrial targeting sequence are also included in Table 1. Class I FHs are sparsely distributed in both uni-and multicellular eukaryotes and are of the single-subunit type. While most multi-cellular eukaryotes with class I FH also have class II FH, *Hymenolepis microstoma* and *Echinococcus multilocularis* (flatworms) are the only multicellular eukaryotes that have only class I *fh* gene. Eukaryotes including *Entamoeba histolytica, Hymenolepis microstoma, Echinococcus granulosus, Gonium pectorale, Chrysochromulina sp.,* and organisms belonging to Alveolata and Kinetoplastida having only class I FH are all parasitic in nature with the exception of *Chrysochromulina sp.,* and *Gonium pectorale* that are free-living. Most other eukaryotes having class I FH also have the gene for class II FH. *Vitrella brassicoformis,* a photosynthetic ancestor of Apicomplexans (25) has genes for both class I and class II type FH, suggesting the occurrence of a gene loss event with respect to class II FH during the evolution of the Apicomplexan lineage as has been noted previously (20). Of special note are organisms belonging to Kinetoplastida that have two genes for class I FH; one encoding the mitochondrial and other the cytosolic enzyme. Upon a search for possible mitochondrial localization of class I and class II FH sequences listed in Table 1 using MitoFates (26), it was seen that in many organisms where both class I and class II FHs are present, class I FH is predicted to localize to mitochondria and not class II FH.

**Table 1.**
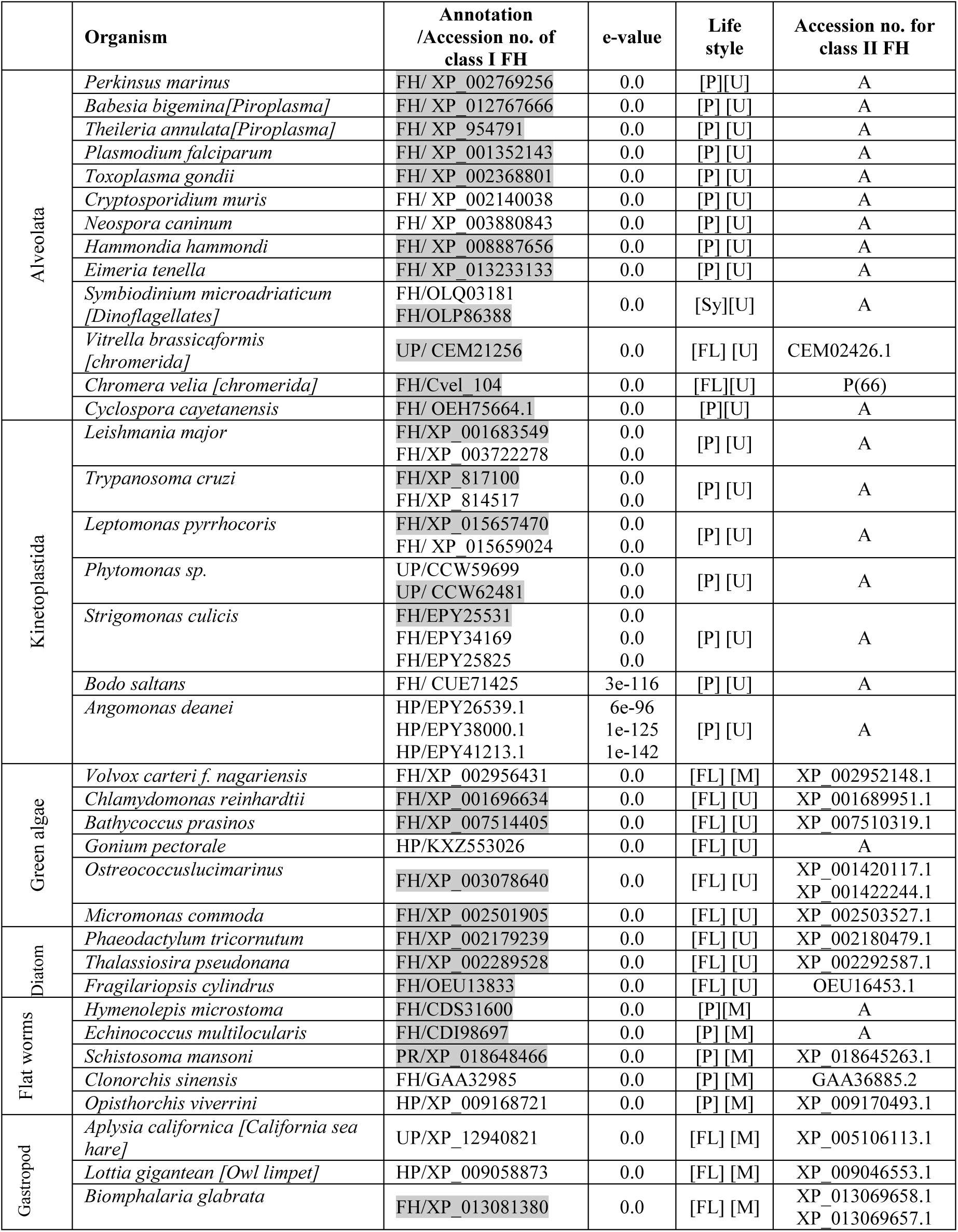
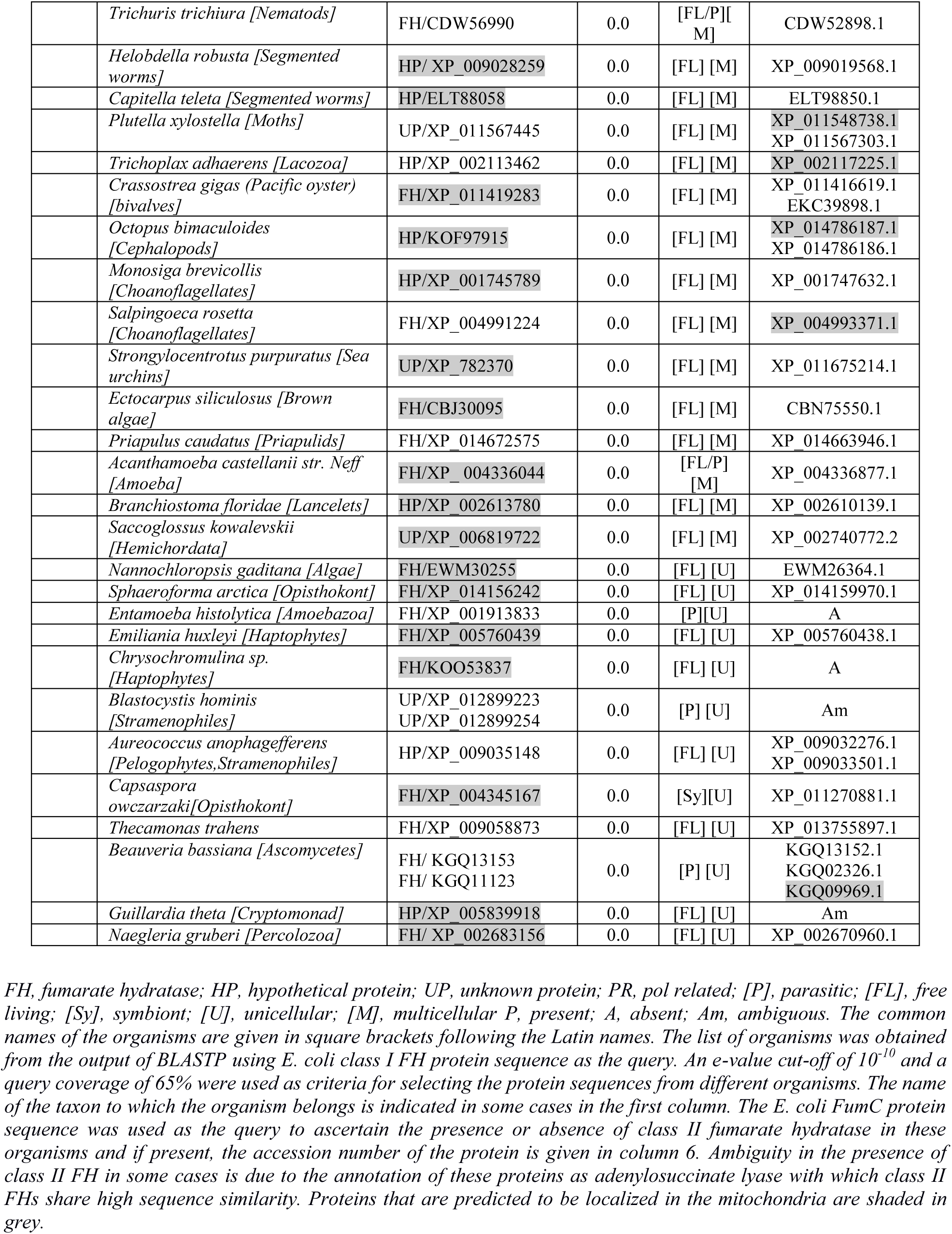
Eukaryotic organisms with class I fumarate hydratase.

### Mitochondrial localization of P. falciparum FH

FH in eukaryotes is known to be localized to mitochondria. Though biochemical evidence suggests that FH is mitochondrially localized in *P. falciparum* (3), microscopic images showing localization to this organelle are not available. To examine the localization of the protein in *P. falciparum,* the *fh* gene on chromosome 9 was replaced with DNA encoding FH-RFA (Fumarate hydratase-regulatable fluorescent affinity tag that comprises GFP, EcDHFR degradation domain and a hemagglutinin tag in tandem) fusion protein by single crossover recombination in PM1KO strain of the parasite (Fig 1a). The genotype of the strain (Fig 1b) was validated by PCR using primers P1-P4 (S1 Table) and used for live-cell imaging after staining with DAPI and MitoTracker Red CM-H_2_XRos. The GFP-positive parasites clearly showed colocalization of GFP signal with MitoTracker Red staining (Fig 1c) showing mitochondrial localization of fumarate hydratase in *P. falciparum*.

**FIGURE 1.**
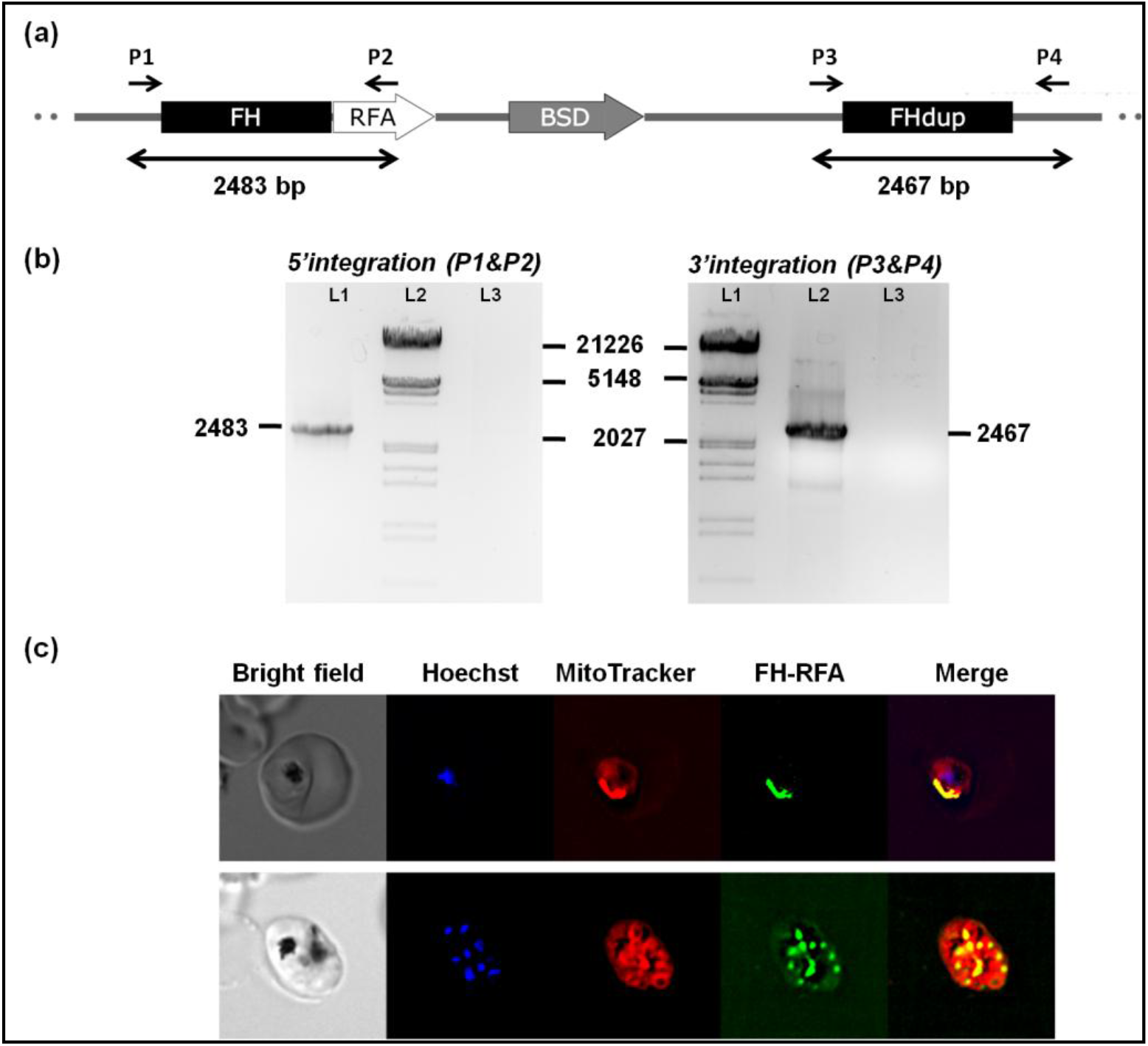
Generation of PfFH-GFP strain encoding FH-GFP and localisation of PfFH. (a) Scheme showing the integration locus with the RFA (GFP+DHFRdd+HA)-tag in tandem with *fh* gene in the strain PfFH-GFP. Oligonucleotides P1 and P2 and, P3 and P4 were used for checking the 5’-and 3’-integration, respectively (S1 Table). P1 and P4 are beyond the sites of integration in the genome. (b) Left panel, genotyping by PCR for validating 5’ integration. The templates used in different lanes are, L1, genomic DNA from PfFH-GFP, L3, *P. falciparum* PM1KO genomic DNA. A band of size 2483 bp validates 5’ integration. Right panel, genotyping by PCR for validating 3’ integration. The templates used in different lanes are, L2, genomic DNA from PfFH-GFP, L3, *P. falciparum* PM1KO genomic DNA. A band of size 2467 bp validates 3’ integration. Molecular weight markers are in lanes L2 and L1 in right and left panels, respectively. (c) Upper panel shows a trophozoite and the lower panel, a schizont. As evident from the merge PfFH localizes to the mitochondrion.

In order to predict the mitochondrial targeting signal in FH sequences from different *Plasmodium* species, different algorithms were used and the results are summarized in S2 Table. Except for *P. knowlesi* FH, wherein all the software were able to predict the signal sequence, none of the FH sequences from other *Plasmodium* species had a conventional targeting signal that could be unambiguously predicted. Of all the mitochondrial proteins predicted by PlasMIT, an artificial neural network based prediction tool developed specifically for predicting mitochondrial transit peptides in *P. falciparum* protein sequences, fumarate hydratase was the only ‘false negative’ (27). This suggests that PfFH localization to the mitochondrion is possibly mediated through an internal signal sequence or an unconventional mitochondrial localization signal.

### PfFH complements fumarase deficiency in E. coli

In order to recombinantly express organellar proteins in *E. coli*, it is preferable to use the DNA sequence corresponding to only the mature protein with the signal peptide deleted. Since none of the bioinformatic prediction tools was able to identify an unambiguous signal sequence in *P. falciparum* FH, we resorted to multiple sequence alignment with bacterial single-subunit type and archaeal two-subunit type FH for generating N-terminal deletion constructs. Examination of the multiple sequence alignment shows a 120 amino acid insertion at the N-terminus in *Plasmodium* FHs that is absent in bacterial and archaeal FH sequences (S1 Fig). Of the N-terminal 120 amino acid residues in *Plasmodial* FHs, the first 40 residues are diverse, while residues 40-120 show a high degree of conservation (S2 Fig) within the genus. Hence, for functional complementation in *E. coli fh* null mutant, three different expression constructs of PfFH protein in pQE30 were generated; that expressing the full length (PfFHFL), N-terminal 40 residues deleted (PfFHΔ40) and N-terminal 120 residues deleted (PfFHΔ120) enzymes.

*E. coli* has three genes that encode fumarate hydratase; *fumA* and *fumB* of the class I type and *fumC* of the class II type. *fumA* and *fumC* genes are in tandem and are driven by a common promoter (7, 28). Starting with JW4083-1, a Δ*fumB* strain of *E. coli,* a triple knockout Δ*fumACB* strain, in which all the three major *fum* genes (*fumA*, *fumC* and *fumB*) are deleted, was generated and validated by PCR (S3 Fig). As expected, while the strain was able to grow normally in malate containing minimal medium (Fig 2a), it was unable to grow on minimal medium containing fumarate as the sole carbon source (Fig 2b). As expected all transformants (containing pQE-PfFHFL, pQE-PfFHΔ40, pQE-PfFHΔ120 and pQE30) of Δ*fumACB* strain of *E. coli* grew well on malate containing minimal medium plates (Fig 2c). In fumarate-containing M9 plates, the cells expressing PfFHΔ40 and PfFHFL grew faster, whereas, the growth rate of cells expressing PfFHΔ120 was slower and no growth of cells carrying just pQE30 was observed (Fig 2d). This shows that the PfFH can functionally complement the deficiency of fumarate hydratase activity in Δ*fumACB* strain and validates that the *P. falciparum* enzyme is indeed fumarate hydratase. The slow growth of PfFHΔ120 expressing Δ*fumACB E. coli* strain indicates that residues 40-120 play a role in the structure and/or function of PfFH despite these residues being conserved only in *Plasmodial* fumarate hydratase sequences and not in others (S3 Fig).

**FIGURE 2.**
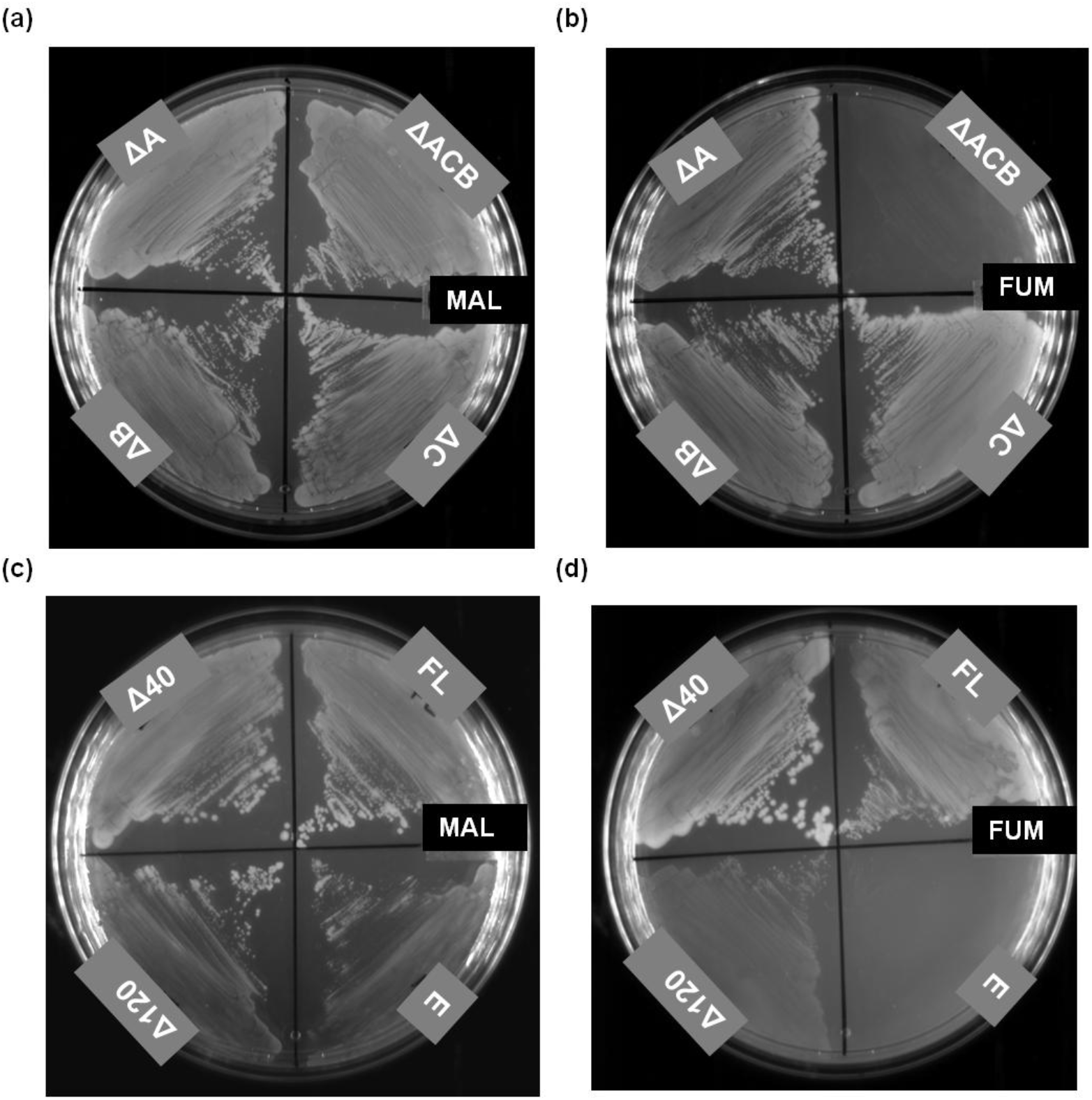
Phenotyping of the *E. coli* strain ΔfumACB and functional complementation by *P. falciparum* FH. Growth phenotype of the *E. coli* strains with at least one copy of fumarate hydratase gene deleted on a) malate and (b) fumarate containing minimal medium. As it is evident from the phenotype, ΔfumACB strain is not able to grow on fumarate containing minimal medium. The growth of ΔfumACB strains expressing either PfFHFL or PfFHΔ40 or PfFHΔ120 of *P. falciparum* fumarate hydratase on (c) malate-and (d) fumarate-containing minimal medium. The plates were scored after 48 h of incubation at 37 °C. ΔfumACB strain containing just pQE30 (E) was used as a control. The experiment was repeated thrice and the images correspond to one of the replicates.

### Activity of PfFHΔ40

PfFΔ40 was expressed with an N-terminal (His)_6_-tag in Codon plus BL21 (DE3) RIL, purified using Ni-NTA affinity chromatography (Fig 3a) and reconstituted *in vitro* with Fe-S cluster. The UV-visible spectrum with absorption maxima at 360 and 405 nm indicates the presence of 4Fe-4S cluster in the enzyme. The addition of sodium dithionite lowered the absorption intensity at 405 nm indicating a reduction of the cluster (Fig 3b) (29–32).The enzyme lacking the reconstituted cluster was devoid of any activity. The activity of PfFΔ40, when examined at 240 nm, showed a time dependent decrease in absorbance with fumarate as the substrate, while with malate an increase was observed. To confirm the chemical identity of the product formed, NMR spectrum was recorded with 2, 3-[^13^C]-fumarate as the substrate. The appearance of two doublets with chemical shift values 70.63, 70.26 ppm and 42.86, 42.29 ppm corresponding to C2, C3 carbons, respectively of malate confirmed that PfFHΔ40 has *in vitro* fumarase activity (Fig 3c).

**FIGURE 3.**
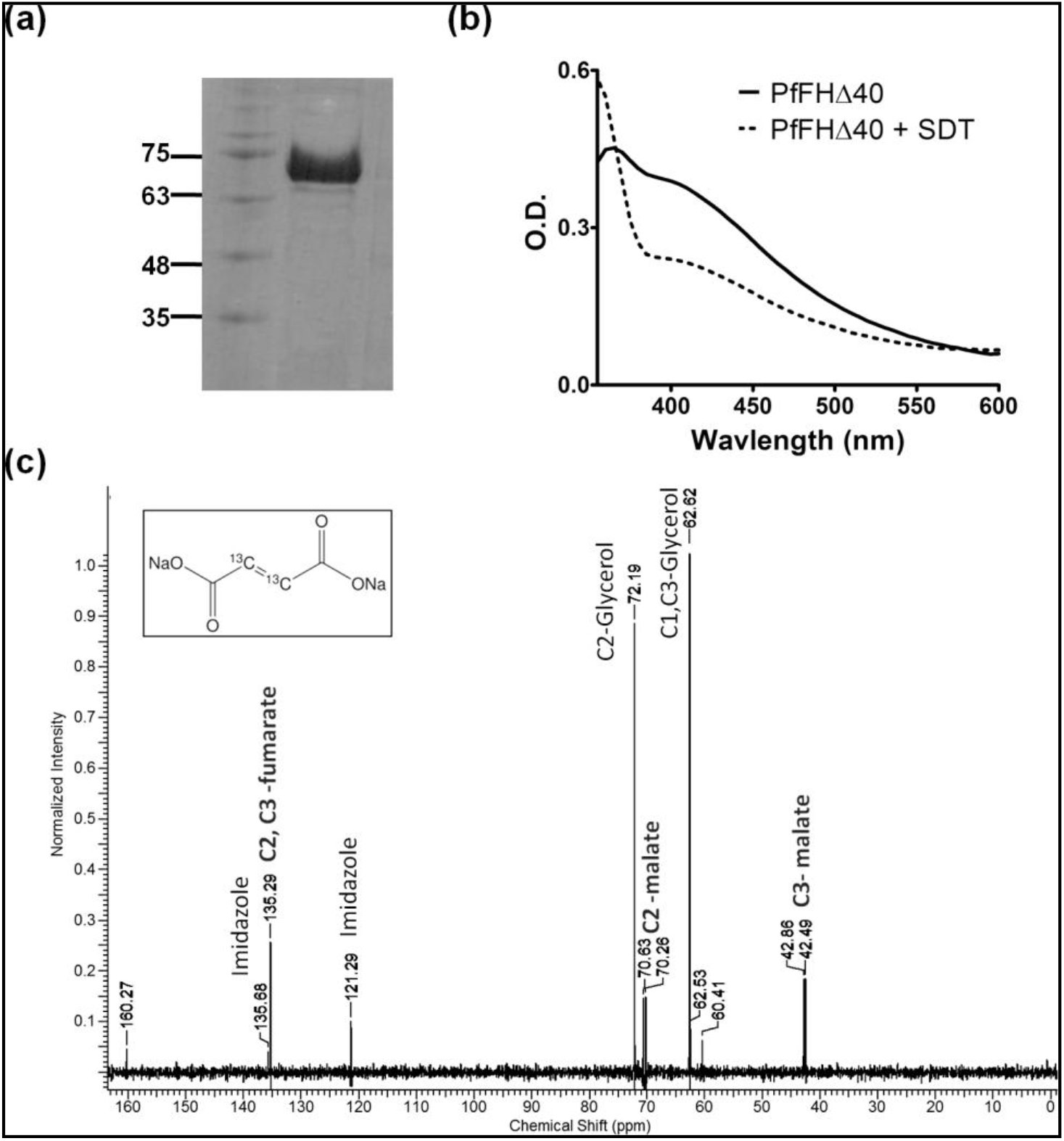
Purification and activity of PfFHΔ40. (a) Lane1, protein molecular weight marker (numbers indicated are in kDa); lane2, Ni-NTA purified PfFHΔ40. (b) The uv-visible absorption spectrum of purified and reconstituted PfFH shows a characteristic peak at 360 and at 405 nm that indicates the presence of a 4Fe-4S cluster. (c) Validation of malate formation by 13C-NMR. The NMR spectrum of assay mixture consisting of 50 μM 2,3-[^13^C]-fumarate in 100 mM potassium phosphate, pH 7.4, incubated with 100 μg of purified PfFHΔ40 enzyme, shows the presence of peaks corresponding to ^13^C-malate. Unreacted ^13^C-fumarate is also present. The inset shows the chemical structure of ^13^C-fumarate. The spectrum is an average of 3000 scans acquired using Bruker 400MHz NMR spectrometer. The peaks corresponding to imidazole and glycerol are from the protein solution

Substrate saturation curves for PfFHΔ40 for both fumarate and malate were hyperbolic in nature indicating the absence of cooperativity. Fit to Michaelis-Menten equation yielded *K*_m_ and *V*_max_ values that are summarized in Table 2. The *K*_m_ values for PfFΔ40 for fumarate and malate in the low millimolar range are similar to that of class I FH from *Leishmania major* (21) and *Trypanosoma cruzi* (22) while for those from bacteria and archaea, the values are in the micromolar range. The catalytic efficiency (*k*_cat_/*K*_m_) of PfFΔ40 is similar to *L. major* FH but 10-100 fold lower than that reported for other class I FHs (Table 2).

**Table 2.** Kinetic parameters of PfFHΔ40 and other class I FH. The units for V_max_, k_cat_ and k_cat_/K_m_ are μmol min^-1^mg^-1^, s^-1^ and s^-1^M^-1^, respectively.

| No | Org./Enzyme name | Substrate | $K_m$ (mM) | $V_{\max}$ or $k_{\text{cat}}$ (#) | $k_{\text{cat}}/K_m$ | Ref. |
| --- | --- | --- | --- | --- | --- | --- |
| 1 | *P. falciparum*<br>*PfFHΔ40* | Fumarate<br>Malate<br>Mesaconate | $2.6 \pm 0.3$<br>$1.2 \pm 0.1$<br>$3.2 \pm 0.3$ | $138 \pm 3$<br>$127 \pm 4$<br>$48 \pm 2$ | $6.2 \times 10^4$<br>$1.3 \times 10^5$<br>$1.7 \times 10^4$ | This study |
| 2 | *T. cruzi*<br>TcFHC<br>TcFHM | Fumarate<br>Malate<br>Fumarate<br>Malate | $0.8 \pm 0.2^*$<br>$2.5 \pm 0.6^*$<br>$1.5 \pm 0.4$<br>$2.8 \pm 0.2$ | $400 \pm 100^\#$<br>$290 \pm 40^\#$<br>$2300 \pm 500^\#$<br>$1050 \pm 40^\#$ | $8.2 \times 10^6$<br>$1.3 \times 10^6$<br>$1.5 \times 10^6$<br>$0.4 \times 10^6$ | (22) |
| 3 | *E. coli*<br>*Fum A*<br>*Fum B* | Fumarate<br>Malate<br>Mesaconate<br>Fumarate<br>Malate<br>Mesaconate | $0.09 \pm 0.02$<br>$0.40 \pm 0.05$<br>$0.22 \pm 0.02$<br>$0.21 \pm 0.03$<br>$0.78 \pm 0.13$<br>$0.10 \pm 0.01$ | $614 \pm 29$<br>$350 \pm 10$<br>$55.6 \pm 1.7$<br>$654 \pm 38$<br>$289 \pm 16$<br>$57.8 \pm 1.4$ | $6.6 \times 10^6$<br>$8.7 \times 10^6$<br>$2.5 \times 10^6$<br>$3.1 \times 10^6$<br>$3.7 \times 10^6$<br>$5.8 \times 10^5$ | (14) |
| 4 | *L. major*<br>*LMFH-1*<br>*LMFH-2* | Fumarate<br>Malate<br>Fumarate<br>Malate | $2.5 \pm 0.4$<br>$2.3 \pm 0.3$<br>$5.7 \pm 1.4$<br>$12.6 \pm 2.7$ | $26.4 \pm 4.4$<br>$11.8 \pm 1.2$<br>$186 \pm 46.5$<br>$138 \pm 18.7$ | $1.1 \times 10^4$<br>$5.6 \times 10^3$<br>$3.6 \times 10^4$<br>$1.2 \times 10^4$ | (21) |
| 5 | *P. furiosus*<br>*FH* | Fumarate<br>Malate | 0.34<br>0.41 | 1376<br>1892 | $3.2 \times 10^6$<br>$3.7 \times 10^6$ | (67) |
| 6 | *P. thermop.*<br>*MmcBC* | Fumarate<br>Malate | 0.43<br>0.59 | 219<sup>#</sup><br>25.2<sup>#</sup> | $5.1 \times 10^5$<br>$4.3 \times 10^4$ | (8) |
| 7 | *B. xerov.*<br>*Bxe_A3136* | Fumarate<br>Malate<br>Mesaconate | $0.10 \pm 0.01$<br>$0.28 \pm 0.02$<br>$0.03 \pm 0.01$ | $296 \pm 5$<br>$118 \pm 3$<br>$117 \pm 6$ | $2.8 \times 10^6$<br>$3.98 \times 10^5$<br>$3.6 \times 10^6$ | (10) |

The substrate promiscuity of PfFHFL, PfFHΔ40, and PfFHΔ120 for other dicarboxylic acids was examined using growth complementation in the *E. coli* strain, Δ*fumACB* (Fig 4). Growth on L-tartrate, D-tartrate, and itaconate was conditional to the presence of PfFH, while growth on meso-tartrate was independent. All three PfFH constructs greatly enhanced the growth of the Δ*fumACB* strain of *E. coli* on mesaconate over the control. *In vitro* activity measurements showed that the parasite enzyme utilizes mesaconate as a substrate converting it to S-citramalate with *K_m_* and *k_cat_/K_m_* values of 3.2 ±0. 3 mM and 1.7 x 10^4^ M^-1^ s^-1^, respectively with the latter value 3.5-and 10-fold lower than that for fumarate and malate, respectively (Table 2). The *in vitro* activity on D-tartrate was measured by a coupled enzyme assay using PfMDH. This activity at 2 mM D-tartrate was 7.8 μmol min^-1^ mg^-1^ that is 9.4-fold lower than that on malate at a similar concentration. The poor growth of Δ*fumACB* strain expressing PfFH constructs on this substrate correlates with the weak *in vitro* activity. Inhibition of PfMDH (the coupling enzyme) at higher concentration of D-tartrate precluded estimation of *k*_cat_ and *K*_m_ values for this substrate. PfFHΔ40 failed to show *in vitro* activity on itaconate (a succinate analog), R-malate and R, R-tartrate (L-tartrate) even at a concentration of 10 mM, indicating that the enzyme is highly stereospecific in recognition of substrates. The growth phenotype of Δ*fumACB* on R R-tartrate and itaconate could arise from PfFH playing a secondary but critical role required for cell growth. These results show that the substrate promiscuity profile of PfFH is similar to class I enzymes from other organisms (7, 9, 14, 33) with the order of preference being fumarate followed by mesaconate and the least preferred being D-tartrate.

**FIGURE 4.**
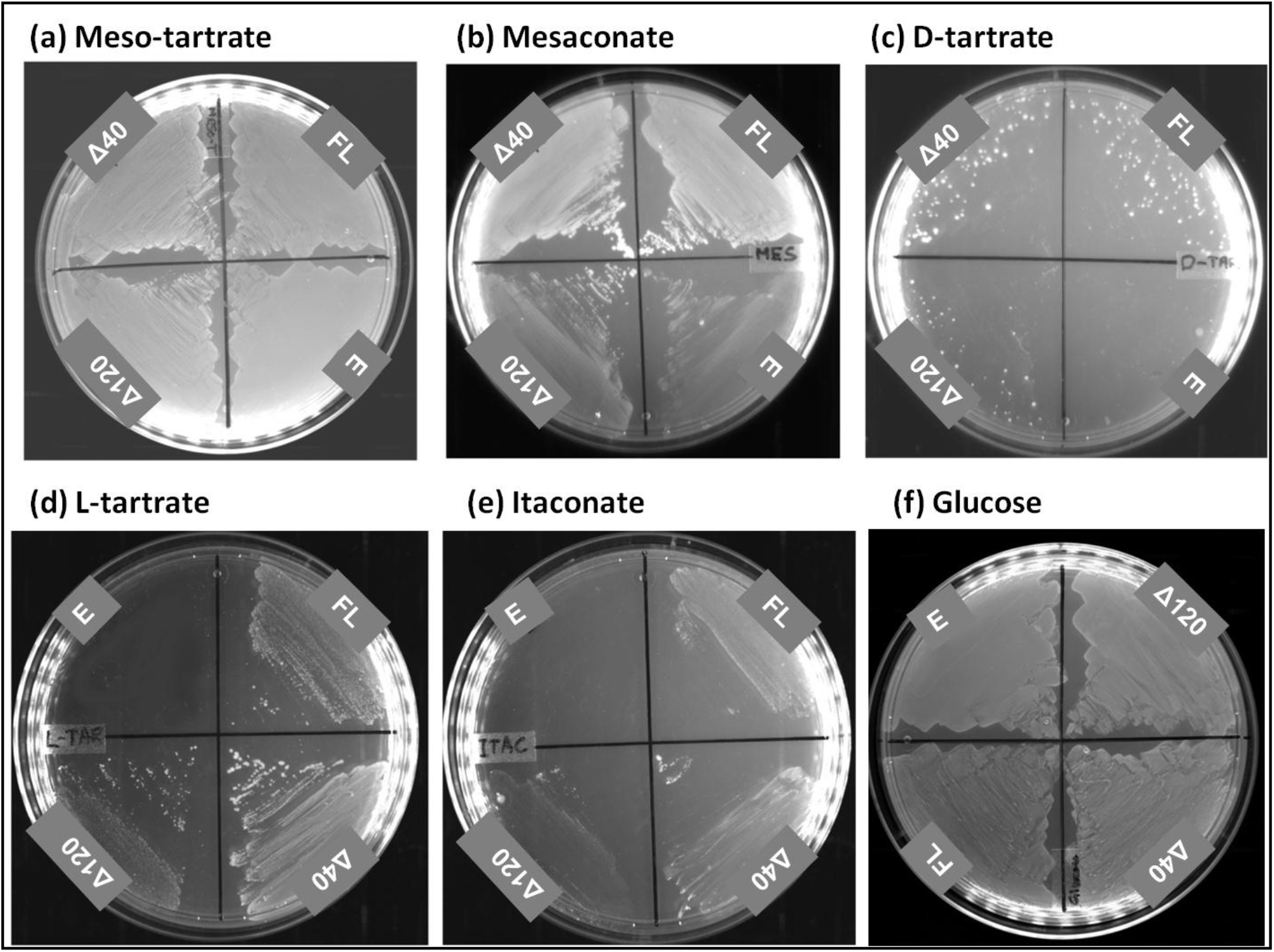
The growth of E.coli strain ΔfumACB expressing PfFH on different carbon sources. The growth of ΔfumACB strains expressing either PfFHFL or PfFHΔ40 or PfFHΔ120 were tested on (a) meso-tartrate, (b) mesaconate, (c) D-tartrate, (d) L-tartrate, (e) itaconate and (f) glucose-containing minimal medium. The plates were scored after 48 h of incubation at 37 °C. ΔfumACB strain containing just pQE30 (E) was used as a control. The experiment was repeated thrice and the images correspond to one of the replicates. The glucose containing plate served as a control for the number of cells plated across the different constructs.

### Mercaptosuccinic acid is class I FH specific inhibitor

Analogs of fumarate, malate, and intermediates of the TCA cycle (including their analogs) were tested for their effect on PfFHΔ40 (S1 text). Of these, the only molecules that inhibited PfFH activity were DL-mercaptosuccinic acid (DL-MSA, Fig 5a) and meso-tartrate. Double reciprocal plots of initial velocity as a function of varied substrate (fumarate and malate) concentrations at different fixed DL-MSA concentrations yielded lines that intersected on the 1/*v* axis indicating the competitive nature of inhibition (Fig 5b and 5c). The K_i_ values for DL-MSA for PfFHΔ40 with malate and fumarate as substrates are 321 ± 26 nM and 548 ± 46 nM, respectively. To test the specificity of MSA for class I FH, its effect on both EcFumA and EcFumC was examined. The Ki value for the inhibition of EcFumA with fumarate as the substrate was 2.9 ± 0.22 μM indicating that DL-MSA is 5.2-fold more potent for PfFHΔ40. It should be noted that the MSA used in the studies is an enantiomeric mixture of DL-isomers and hence, the Ki value would be half of that determined. Further, as D-malate is not an inhibitor of PfFH, only L-MSA would be expected to bind to the enzyme. DL-MSA was also found to inhibit the two-subunit class I FH from *Methanocaldococcus jannaschii* (unpublished) while, there was no effect on EcFumC even at a concentration of 10 mM indicating it’s exclusive specificity for class I FH. A recent study has reported the inhibition of TcFH by DL-MSA with a K_i_ of 4.2 ± 0.5 μM while no effect was observed on the class II human FH (22). Interestingly, DL-MSA is not a substrate for PfFH as seen by spectrophotometric assays at 240 nm with 10 mM DL-MSA and 1 μM enzyme that failed to show either formation of the enediolate intermediate or the product fumarate through the liberation of H_2_S. The reason for DL-MSA’s high specificity for class I FH must stem from the presence of 4Fe-4S cluster that interacts with the C2-hydroxyl group of malate (11). Replacement of the hydroxyl group with a thiol probably leads to tight binding through Fe-S interaction. Also, the formation of a stable DL-MSA-FH complex could arise from the C3-hydrogen of DL-MSA being less acidic than that of malate. The 4Fe-4S cluster containing quinolinate synthase (NadA) is strongly inhibited by dithiohydroxyphthalic acid (DTHPA), the thio-analog of the transition state intermediate of the reaction catalyzed, in a manner similar to MSA inhibition of class I FH. Interactions of the thiol groups of DTHPA with Fe atom of the cluster leads to a strong binding affinity for the enzyme (34). In this context, we expect thiomesaconate and S, S-dithiotartrate, analogs of the substrates mesaconate and S, S, tartrate (D-tartrate) to be also strong inhibitors of class I FH.

**FIGURE 5.**
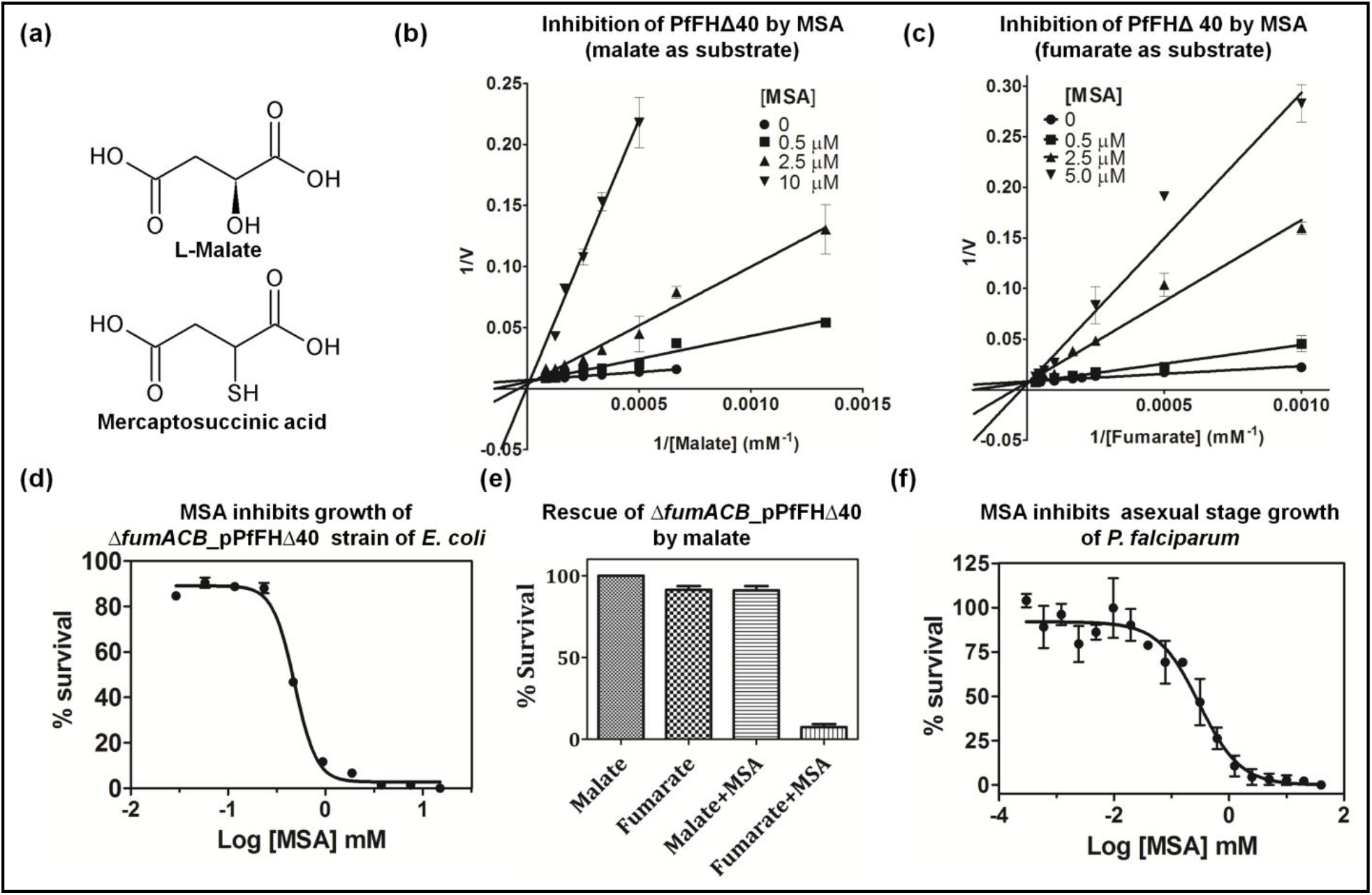
Specificity of DL-MSA for class I FH. (a) Structures of L-malate and mercaptosuccinic acid. (b) Lineweaver-Burk plot of initial velocity at varied malate and different fixed MSA concentrations. (c) Lineweaver-Burk plot of the initial velocity at varied fumarate and different fixed MSA concentrations. (d) Inhibition of the growth of the ΔfumACB_pPfFHΔ40 strain of *E. coli* by MSA. (e) Rescue of MSA mediated growth inhibition of ΔfumACB_pPfFHΔ40 upon addition of malate. (f) Inhibition of the in vitro growth of intraerythrocytic asexual stages of *P. falciparum* by MSA.

Albeit slightly less effective as an inhibitor, meso-tartrate competitively inhibited PfFHΔ40 with a K_i_ value of 114 ± 17 μM. This compound inhibited both EcFumA and EcFumC with similar K_i_ values of 625 and 652 μM, respectively. Meso-tartrate has two chiral carbons with S-configuration at C2, and R-configuration in C3. It should be noted that in the case of both EcFumC and EcFumA S, S-tartrate (D-tartrate) is a substrate (14). The inhibition by meso-tartrate of both class I and II FH indicates relaxed stereospecificity of these enzymes at the C3 carbon of the substrates. Pyromellitic acid, a known potent inhibitor of class II FH had no effect on the activity of the two class I enzymes tested (PfFHΔ40 and EcFumA), while completely abolishing the activity of the class II enzyme (EcFumC). The specificity exhibited by DL-MSA for class I FH and, pyromellitic acid and S-2,3-dicarboxyaziridine (35) for class II FH supports the presence of different active site environments in the two classes of enzymes. This provides a framework for developing class I PfFH (and in general for class I FH) specific inhibitors that will have no effect on the human enzyme.

### Growth inhibition by DL-MSA

Since the growth of *E. coli* Δ*fumACB* strain on minimal medium containing fumarxamined in *P. berghei* with BALB/c mice as host. For this, *fh* gene knockout construct generated through recombineering based strategy was used (S4 Fig). Transfected parasites were injected into mice, selected on pyrimethamine and drug resistant parasites that appeared 10 days after infection were subjected to limiting dilution cloning. All the 17 *P. berghei* clones (A to Q) obtained by limited dilution cloning of the drug resistant parasites were examined by PCR to confirm the presence of the integration cassette and the absence of the *fh* gene. Oligonucleotides used for genotyping of the clones are provided in S1 Table. The expected genomic locus upon the integration of selectable marker cassette by double crossover recombination is shown schematically in Fig 6 a and the wild-type (with *fh* gene) is shown in Fig 6 b. Genotyping by PCR was performed to confirm integration of selectable marker cassette at expected locus (Fig 6 c & as the sole carbon source is conditional to the presence of functional FH, the effect of DL-MSA on the growth of Δ*fumACB*_pPfFΔ40 was examined. DL-MSA inhibited the growth of Δ*fumACB*_pPfFΔ40 with an IC_50_ of 482 ± 4 μM (Fig 5d) and the addition of malate completely rescued the inhibition (Fig 5e). This shows that the toxicity of DL-MSA is indeed due to inhibition of the metabolic conversion of fumarate to malate. Δ*fumACB E. coli* strain can serve as a facile primary screening system for small molecules acting as inhibitors of PfFH as it circumvents *in vitro* assays with the oxygen-sensitive labile enzyme. With DL-MSA as an inhibitor of PfFH under *in vitro* and *in vivo* conditions, the molecule was checked for its toxicity on intraerythrocytic stages of *P. falciparum* in *in vitro* culture. DL-MSA was found to kill parasites in culture with an IC_50_ value of 281 ± 68 μM (Fig 5f). Though DL-MSA is a potent inhibitor of PfFHΔ40 with a K_i_ value of 547 ± 47 nM (with fumarate as substrate), the IC_50_ values for the inhibition of both Δ/umACB_pPfFΔ40 and *P. falciparum* are significantly higher.

### Essentiality of fumarate hydratase for P. berghei is host strain dependent

Earlier attempt at knockout of fumarate hydratase gene in *P. falciparum* was not successful (23). Therefore, the essentiality of FH was examined in *P. berghei* with BALB/c mice as host. For this, *fh* gene knockout construct generated through recombineering based strategy was used (S4 Fig). Transfected parasites were injected into mice, selected on pyrimethamine and drug resistant parasites that appeared 10 days after infection were subjected to limiting dilution cloning. All the 17 *P. berghei* clones (A to Q) obtained by limited dilution cloning of the drug resistant parasites were examined by PCR to confirm the presence of the integration cassette and the absence of the *fh* gene. Oligonucleotides used for genotyping of the clones are provided in S1 Table. The expected genomic locus upon the integration of selectable marker cassette by double crossover recombination is shown schematically in Fig 6 a and the wild-type (with *fh* gene) is shown in Fig 6 b. Genotyping by PCR was performed to confirm integration of selectable marker cassette at expected locus (Fig 6 c & d), presence/absence of the *fh* gene (Fig 6 e) and the presence of the selection cassette (Fig 6 f & g). The results of the PCRs showed that though all the parasite clones carried the selectable marker, hDHFR-yFCU cassette in the genomic DNA (Fig 6 f), they also retained the *fh* gene (Fig 6 e). In two of the clones (C and O), the integration of the cassette was at a random site as they failed to answer for both 5’ and 3’ integration PCRs. 12 clones yielded the expected PCR amplified fragment for 5’ integration (Fig 6 c) while a band of the expected size was not obtained for 3’ integration PCR. One clone (M) yielded expected PCR amplified fragment for only 3’ integration (Fig 6 d) and not for 5’ integration. Integration of the selection cassette through single crossover recombination using 5’ or 3’ homology arm with the intact *fh* gene present downstream or upstream, respectively would yield this PCR result. As the DNA used for transfection was linear, circularization of the fragment must have enabled this single crossover recombination. Only 2 clones (J and Q) answered positive for both 5’ and 3’ integration PCRs while continuing to harbor *fh.* These two clones must have arisen from a double crossover recombination event in a population of parasites harboring the duplicated copy of *fh.* Although parasites with gene duplication are thought to be unstable, the existence of duplication has been noted earlier in the case of *rio2* (36) and *dhodh* (37). The variation in the genotype across the 17 clones that we have obtained shows that the parasites have not multiplied from a single wrong event of homologous recombination. On the contrary, the clonal lines with different genotypes, continuing to harbor *fh* suggests a strong selection pressure for the retention of this gene. The PCRs with primers P9 and P10 (Fig 6 e) encompassing the full-length gene yielded the expected size band with genomic DNA from all 17 clones, indicating that all clones contain full-length *fh* gene.

**FIGURE 6.**
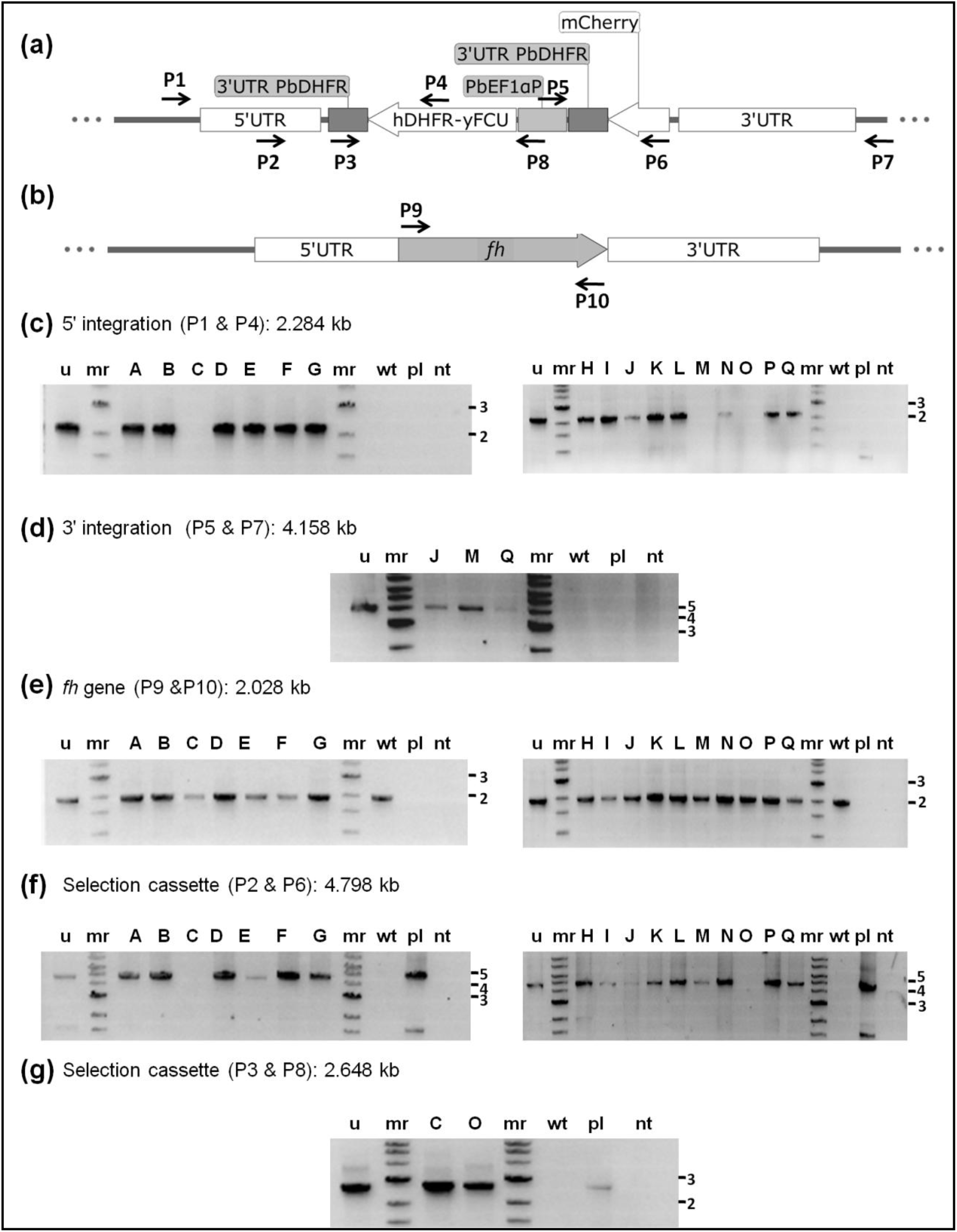
Genotyping of *P. berghei* clones of knockout of fumarate hydratase. (a) Schematic representation of the selectable marker cassette inserted into the fh gene locus of *P. berghei* genome. Primers (P1-P8) used for diagnostic PCRs are indicated. (b) Schematic representation of the fh gene (PBANKA_0828100) flanked by 5’ UTR and 3’ UTR showing the location of primers P9 and P10. (c) clones A-G (left panel) and clones H-Q (right panel) for detection of 5’ integration; (d) clones J, M and Q for detection of 3’ integration (other clones did not answer for this PCR); (e) clones A-G (left panel) and clones H-Q (right panel) for the detection of fh gene; (f) clones A-G (left panel) and H-Q (right panel) for the presence of selectable marker cassette; (g) clones C and O using primers P3 and P8. Clones C, M and O did not answer for 5’ integration while only clones J, M and Q answered for 3’ integration. All clones answered for the presence of the fh gene (Panel d). All clones except C and O answered by PCR with primers P2 and P6 indicating the integration of the entire selectable marker cassette into the genome. Clones C and O answered for a shorter fragment of the selectable marker cassette covered by primers P3 and P8. hDHFR-yFCU, human DHFR-yeast cytosine and uridyl phosphoribosyltransferase, u, uncloned population, mr, molecular weight marker; wt, wild-type *P. berghei* genomic DNA; pl, pJAZZ-FH knockout construct (supplementary figure S2); nt, control PCR without template. Numbers to the right of panels d, e, f, g and h are the sizes of the marker DNA fragments in kbp.

A study that appeared recently reports on the knockout of *fh* gene in *P. berghei,* with the knockout parasites exhibiting slow growth phenotype (38). Though the authors show the absence of *fh* gene expression in the knockout strain, the genotyping for confirmation of knockout that was carried out with oligonucleotide primers corresponding to the homology arm used for recombination cannot confirm the site of integration. Apart from the length of the homology arm used for recombination, the key difference is with regard to the strains of mice used. While our study has used BALB/c, Niikura *et al.,* (38) have used the C57BL/6 strain of mice. *P. berghei* is known to exhibit differences in growth and infectivity across different strains of mice (39–41). This prompted us to examine the essentiality of *fh* gene for *P. berghei* when grown in the two different mouse strains, C57BL/6 and BALB/c. For this, a single transfection mixture was split into two halves and injected into C57BL/6 and BALB/c mice and the whole experiment was performed twice. In the two experiments, intravenous injection of the transfected *P. berghei* cells yielded parasites in both strains of mice, C57BL/6 and BALB/c. However, in one of the attempts, upon pyrimethamine selection, drug resistant parasites appeared only in the C57BL/6 mouse and not in the BALB/c strain even after 20 days of observation. In the second attempt, pyrimethamine resistant parasites were obtained in both C57BL/6 and BALB/c mice. Genotyping by PCR of drug selected parasites using diagnostic oligonucleotides was performed. Drug selected parasites obtained from C57BL/6 mice of both attempts of transfection showed the right integration of marker cassette and along with the absence of *fh* gene (results of genotyping performed from the second attempt of transfection is shown in Fig 7). On the contrary, genotyping by PCR of drug selected parasites obtained from BALB/c mouse, revealed the presence of the marker cassette (Fig 7b) along with the *fh* gene (Fig 7a). Results from the transfection experiments, taken together, indicate that *fh* gene in *P. berghei* can be knocked out when the parasites are grown in C57BL/6 strain of mice and not when BALB/c mouse is the host. Similar mouse-strain specific essentiality of a *P. berghei* gene is seen in the case of purine nucleoside phosphorylase (PNP). While *P. berghei* PNP has been shown to be refractory to knockout in transfectants grown in BALB/c mice as deposited in PhenoPlasm database by Sanderson and Rayner (42, 43), Niikura *et al,* have successfully deleted the gene in *P. berghei* when transfected and grown in C57BL/6 mice (44). It should be noted that the exact nature of the plasmid constructs used for knockout are different across the two studies. To our knowledge, the study reported here is the first where simultaneously the same knockout-construct has been used for deletion of *fh* gene using two different strains of mice as hosts. The variation that we observe across the two hosts used suggests the role of mouse-strain in determining the essentiality of a parasite gene.

**FIGURE 7.**
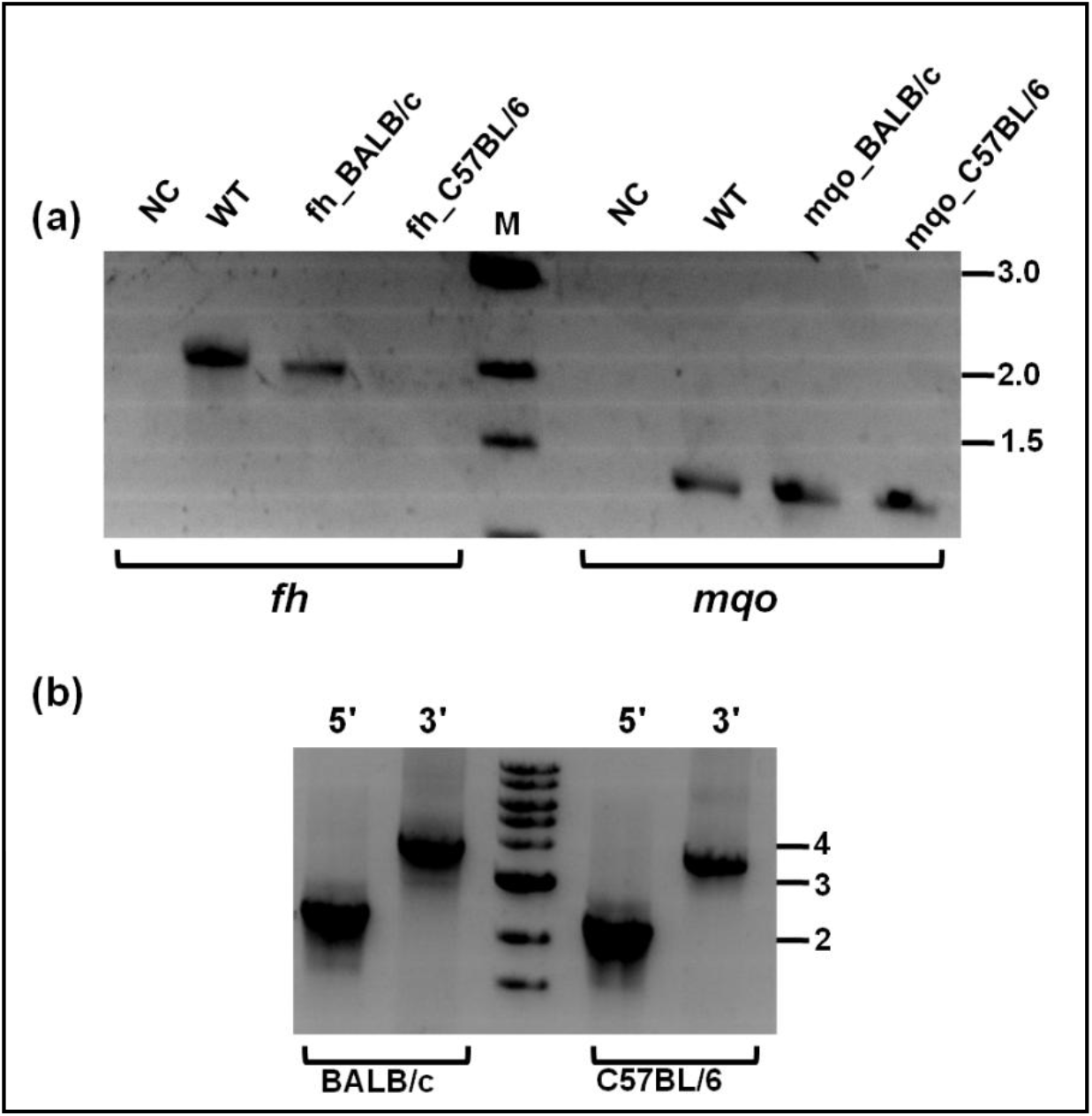
Genotyping of transfectants grown and drug-selected in different mice strain. (a) The presence/absence of fumarate hydratase gene was validated by PCR using oligonucleotides P9 and P10 and genomic DNA isolated from respective parasites as template (lanes bracketed as fh). As a positive control for the presence of genomic DNA, PCR was performed with oligonucleotides corresponding to a segment of *mqo* gene loci (lanes bracketed as *mqo*). (b) Validation of 5’-and 3’-integration using primer pairs P1 and P4 and, P5 and P7, respectively. NC, negative control lacking template DNA, WT, *P. berghei* ANKA wild-type genomic DNA, BALB/c, genomic DNA from transfectants grown in BALB/c mouse, C57BL/6, genomic DNA from transfectants grown in C57BL/6 mouse. The sequences of the oligonucleotides used are shown in supplementary table 1.

## CONCLUSION

Unlike higher eukaryotes including humans that have only class II fumarate hydratase, most parasitic protozoa have only the class I enzyme. The class I FH in *P. falciparum* localizes only to the mitochondrion, though a mitochondrial targeting signal sequence could not be identified using any of the bioinformatic tools. Upon expression of recombinant proteins in *E. coli,* of the three constructs (PfFHFL, PfFHΔ40, and PfFHΔ120) used, the highest level of soluble protein was obtained with PfFHΔ40. The purified, Fe-S cluster-reconstituted PfFHΔ40 was active and catalyzed the reversible conversion of fumarate and malate with a lower *K*_m_ value for malate and similar *k*_cat_ values for both the forward and reverse reactions. The parasite enzyme complements fumarase deficiency in *E. coli.* Both *in vitro* and *in vivo* in *E. coli,* PfFH exhibits extended substrate specificity for D-tartrate and mesaconate. The significance of the extended substrate specificity of PfFH for mesaconate and D-tartrate with regard to *Plasmodium* cellular biochemistry is unclear at this stage.

MSA is a highly potent and specific inhibitor of class I FH *in vitro.* Surprisingly, *in vivo*, in both *E. coli* expressing PfFH and intraerythrocytic *P. falciparum*, the IC_50_ values for cell death are significantly higher (in the micromolar range). The rescue of the growth inhibition of the *E. coli* cells by malate indicates that the inhibitory effect *in vivo* is specifically through PfFH. The discrepancy between the K_i_ value for the purified enzyme and the IC_50_ value on cells could be either due to the low intracellular availability of the drug (owing to the hydrophilic nature of the molecule and hence poor transport) or due to metabolism leading to degradation of MSA. MSA dioxygenase, an enzyme that converts mercaptosuccinic acid to succinate is present in the bacterium *Variovorax paradoxus* (45, 46). We, however, could not find homologues of the enzyme in *E. coli* or *P. falciparum.* In addition, in *P. falciparum*, the host erythrocyte in which the parasite resides contains the class II enzyme that is not inhibited by MSA and this could substitute for the lack of parasite FH activity due to inhibition. However, under these conditions, regeneration of aspartate (3) and NAD^+^ pools from fumarate mediated by PfFH would require the transport of intermediates across the 3 compartments (erythrocyte, parasite and the mitochondrion), evidence for which is absent. Therefore, the effectiveness of substitution of PfFH by the human enzyme is unclear.

Attempts at knockdown of PfFH levels in the FH-GFP strain where FH is fused to EcDHFR degradation domain did not result in lowering of protein levels. It has been shown that proteins targeted to organelles and possessing a signal sequence lack the accessibility to proteasomal degradation machinery and hence the conditional degradation approach may be unsuitable for achieving knockdown of protein levels (47). Knockout of *fh* in *P. berghei* using BALB/c strain of mice yielded parasites that showed insertion of the marker cassette with the retention of the functional copy of the gene suggesting its essentiality for the intra-erythrocytic stages. The major source of intracellular fumarate in *Plasmodium* is from the synthesis of AMP. From the context of metabolism, the absence of fumarate hydratase would result in the accumulation of fumarate. The possible metabolic consequences of this are schematically shown in Fig 8. The last reaction in AMP synthesis catalyzed by ASL is a reversible process with similar catalytic efficiencies for the forward and reverse reactions. Accumulation of fumarate could lead to an increased flux through the reverse reaction catalyzed by ASL resulting in accumulation of sAMP and lowered levels of AMP, eventually resulting in compromised cell growth. Subversion of ASL activity through the use of AICAR has been shown to result in parasite death (48). Apart from perturbing AMP synthesis, high levels of fumarate can result in succination of cysteinyl residues in proteins and glutathione (49) thereby, compromising cellular homeostasis (50). Succinated proteome in human cell lines (51–53) and *Mycobacterium tuberculosis* (54) have been examined and these studies highlight the toxic effects of high levels of fumarate. Fumarate is recycled to aspartate through the action of enzymes, FH, MQO and AAT. In the absence of FH, the levels of malate and oxaloacetate intermediates in this pathway would be perturbed leading to lower levels of recycling. Further, with lowered levels of OAA due to the absence of FH, the generation of NAD^+^ through MDH would also be impaired due to the absence of FH. All these biochemical requirements could make FH in *Plasmodium* essential. Identification of host-specific factors responsible for the differential essentiality of *fh* gene in *P. berghei* might help elucidate the biochemical need of fumarate hydratase in the human malaria parasite, *P. falciparum.*

**FIGURE 8.**
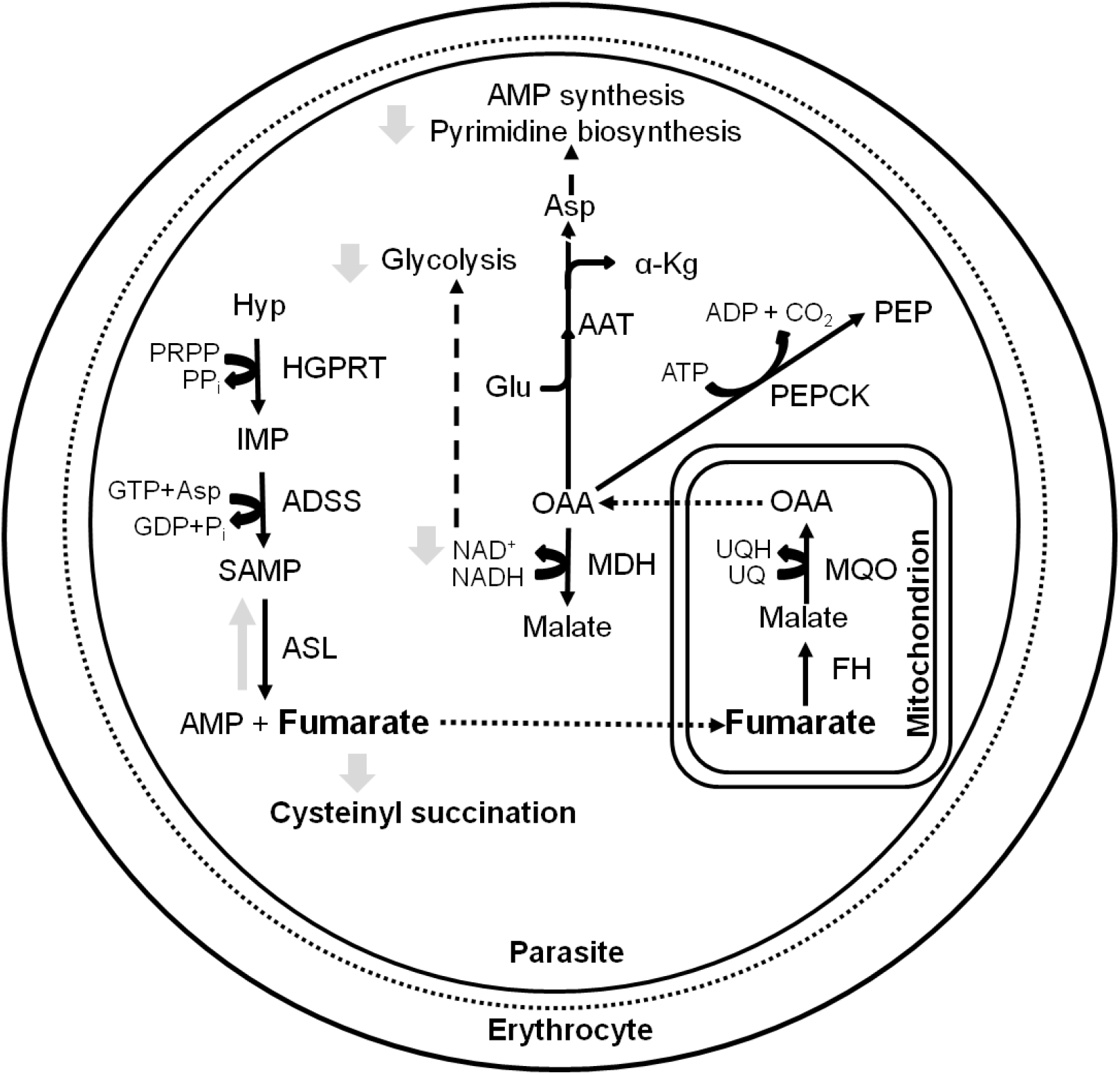
The metabolic consequences of fumarate hydratase gene deletion in *Plasmodium.* Dashed arrows indicate the flow of metabolites into a pathway while dotted arrows indicate transport across compartments. Grey arrows show possible metabolic consequences of fh gene deletion. AAT, aspartate aminotransferase, ADSS, adenylosuccinate synthetase, ASL, adenylosuccinate lyase, FH, fumarate hydratase, HGPRT, hypoxanthine-guanine phosphoribosyltransferase, MDH, malate dehydrogenase, MQO, malate-quinone oxidoreductase, PEPCK, phosphoenolpyruvate carboxykinase, α-Kg, α-ketoglutarate, AMP, adenosine 5’-monophosphate, Asp, aspartic acid, Glu, glutamic acid, Hyp, hypoxanthine, OAA, oxaloacetate, PEP, phosphoenolpyruvate, PRPP, phosphoribosyl 5’-pyrophosphate, SAMP, succinyl-AMP, UQ, ubiquinone, UQH, ubiquinol.

## EXPERIMENTAL PROCEDURES

### Materials

RPMI-1640, components of cytomix and all chemical reagents used were obtained from Sigma Aldrich., USA. MitoTracker Red CM-H_2_XRos, Hoechst 33342, AlbuMAX I, Ni-NTA conjugated agarose and Phusion high-fidelity DNA polymerase were procured from Thermo Fisher Scientific Inc., USA. Restriction enzymes and T4 DNA ligase were from New England Biolabs, USA. Primers were custom synthesized from Sigma-Aldrich, Bangalore. Media components were from Himedia Laboratories, Mumbai, India. 2, 3-[^13^C]-fumarate was procured from Isotec, Sigma Aldrich, USA and DL-mercaptosuccinic acid (DL-MSA) was obtained from Sigma Aldrich, USA.

### Sequence analysis-E. coli

FumA (EcFumA) protein sequence (UniProt ID: P0AC33) was used as a query in BLASTP (55) to retrieve all eukaryotic class I FH sequences by restricting the search to eukaryotes. Each of these eukaryotic organisms with class I FH was individually searched for the presence of class II FH using BLASTP and *E. coli* FumC (EcFumC) (UniProt ID: P05042) as the query sequence. Hits with an e-value lower than10^-10^ were considered significant.

### Generation of plasmid constructs

Sequences of all oligonucleotide primers used for cloning and for genotyping of *E. coli* and *Plasmodium* mutants are given in S1 Table. For recombinant-expression of PfFH, the DNA fragment corresponding to a protein segment lacking the N-terminal 40 amino acids (Δ40) was amplified by PCR using parasite genomic DNA as template, appropriate oligonucleotides and Phusion DNA polymerase. The fragment was cloned into modified pET21b (Novagen, Merck, USA) using restriction enzyme sites BamHI and SalI, to obtain the construct pET-PfFHΔ40 that encodes the protein with an N-terminal (His)_6_-tag. For functional complementation in the *fh* null strain of *E. coli,* pQE30 plasmid (Qiagen, Germany) containing full length (PfFHFL) and two different N-terminus deleted constructs (PfFHΔ40, PfFHΔ120) of the *P. falciparum fh* gene were used. The generation of different expression constructs involved amplification by PCR of appropriate fragments followed by cloning into pQE30 plasmid using restriction sites BamHI and SalI. The plasmids thus obtained are pQE-PfFHFL, pQE-PfFΔ40 and pQE-PfFΔ120. For 3’-tagging of the endogenous *fh* gene with GFP in the *P. falciparum* strain PM1KO (56), the nucleotide fragment corresponding to the full-length *fh* gene without the terminator codon was amplified from *P. falciparum* 3D7 genomic DNA using appropriate oligonucleotides (PfFHpGDB-Xho1-FP and PfFHpGDB-AvrII-RP; S1 Table) and cloned into the plasmid, pGDB (56) using the restriction sites XhoI and AvrII. This plasmid construct is referred to as pGDB-PfFH. For recombinant expression of *E. coli* FumC and FumA enzymes, the nucleotide sequence corresponding to the full-length genes were PCR amplified using *E. coli* genomic DNA as template and cloned in pQE30 and pET-DUET (Novagen, Merck), respectively using the restriction sites BamHI and SalI. The resulting plasmids are pQE-EcFumC and pET-EcFumA. All the clones were confirmed by DNA sequencing.

### Protein expression, purification and reconstitution of iron-sulfur cluster

For recombinant expression of PfFΔ40 and EcFumA, the *E. coli* strain BL21(DE3)-RIL was transformed with pET-PfFΔ40/ pET-EcFumA and selected on Luria-Bertani agar (LB agar) plate containing ampicillin (100 μg ml^-1^) and chloramphenicol (34 μg ml^-1^). Multiple colonies were picked and inoculated into 10 ml of LB broth. The culture was grown for 6 h at 37 °C, the cells were pelleted, washed with antibiotic free LB broth and then used for inoculating 800 ml of Terrific broth (TB). The cells were grown at 30 °C until OD_600_ reached 0.5, thereafter induced with IPTG (0.05 mM for PfFHΔ40 and 0.3 mM for EcFumA) and grown for further 16 h at 16 °C for PfFHΔ40 and 4 h at 30 °C for EcFumA. Cells were harvested by centrifugation and resuspended in lysis buffer containing 50 mM Tris HCl, pH 7.4, 150 mM NaCl, 1 mM PMSF, 5 mM β-mercaptoethanol and 10 % glycerol. Cell lysis was achieved by 4 cycles of French press at 1000 psi and the lysate cleared by centrifugation at 30,000 x g for 30 minutes. The supernatant was mixed with 1 ml of Ni-NTA agarose slurry pre-equilibrated with lysis buffer and incubated in an anaerobic chamber for 30 minutes at room temperature. The tube was sealed airtight within the chamber and transferred to 4 °C. Binding of the (His)_6_-tagged PfFHΔ40/EcFumA to Ni-NTA agarose was continued with gentle shaking for 3 h. The tube was transferred back to the chamber and the beads were washed with 50 ml of lysis buffer followed by washes with 10 mM and 20 mM imidazole containing lysis buffer (10 ml each) and the protein eluted directly with 500 mM imidazole in lysis buffer. An equal volume of 100 % glycerol was added to the eluate such that the final concentration of glycerol is 50 %. EcFumC was purified in aerobic conditions using Ni-NTA affinity chromatography.

Reconstitution of the cluster in PfFHΔ40 and EcFumA was performed under anaerobic conditions. For PfFHΔ40, the procedure was initiated by incubation of the protein solution with 5 mM DTT for 30 min. Following this, 0.5 mM each of sodium sulfide and ferrous ammonium sulfate was added. The reconstitution was allowed to proceed overnight following which the protein was used for activity measurements. For EcFumA, reconstitution was achieved by the addition of 5 mM DTT for 30 min followed by the addition of 0. 5 mM ferrous ammonium sulfate.

### Activity measurements

For recording NMR spectra, the purified recombinant PfFHΔ40 was incubated with 2, 3-[^13^C]-fumarate for 30 min at 37 °C in 20 mM sodium phosphate, pH 7.4. The protein was precipitated with TCA and the supernatant, neutralized with 5 N KOH was used for recording ^13^C-NMR spectrum in a 400 MHz Agilent NMR machine. D_2_O was added to a final concentration of 10 % to the sample before acquiring the spectrum.

All initial velocity measurements were performed at 37 °C using a spectrophotometric method and initiated with the addition of the enzyme. The activity of EcFumC was measured using a reported method (11) in a solution containing 100 mM MOPS, pH 6.9, 5 mM MgCl2, and 5 mM DTT. For EcFumA, the assays were performed in 50 mM potassium phosphate, pH 7.4 containing 2 mM DTT. The activity of PfFΔ40 was found to be maximal at pH 8.5 and all assays were carried out at this pH in 50 mM Tris-HCl. The conversion of fumarate to malate was monitored spectrophotometrically as a drop in absorbance caused by the depletion of fumarate. Depending upon the initial concentration of fumarate, the enzymatic conversion was monitored at different wavelengths; 240 nm (ε_240_ = 2440 M^-1^ cm^-1^) (9) for fumarate concentrations of up to 500 μM, 270 nm (ε_270_ = 463 M^-1^ cm^-1^) for concentrations ranging from 0.5 to 1.2 mM, 280 nm (ε_280_ = 257 M^-1^ cm^-1^) for concentrations from 1.2 to 2.6 mM, 290nm (ε_290_ = 110 M^-1^ cm^-1^) for concentrations from 2.6 to 6 mM, 300 nm (ε_300_ = 33 M^-1^ cm^-1^) for concentrations from (6 to 20 mM) and 305 nm (ε_305_= 18 M^-1^ cm^-1^) for concentrations from (20 to 40 mM) in a quartz cuvette of 1 cm path length. The conversion of mesaconate to citramalate was monitored as drop in absorbance at wavelengths 240 nm (ε_240_ = 3791 M^-1^ cm^-1^) for concentrations up to 250 μM, 280 nm (ε_280_ = 142 M^-1^ cm^-1^) for concentrations from 250 μM to 4000 μM, and at 290 nm (ε_290_ = 40 M^-1^ cm^-1^) for concentrations from 4-16 mM. The use of different wavelengths ensured that the sensitivity of the detection of conversion of fumarate to malate was maximal. The conversion of malate to fumarate was monitored spectrophotometrically as an increase in absorbance at 240 nm due to the synthesis of fumarate. Activity on tartrate was monitored by a coupled enzyme assay using *P. falciparum* malate dehydrogenase (PfMDH) purified in-house from an *E. coli* expression clone (3). The assay was carried out at 37 °C in 50 mM Tris HCl, pH 8.5 containing 100 μM NADH, 4 μg PfMDH, and 2 mM D-tartrate. The reaction was initiated with 3.4 μg of PfFH.

For testing the effect of small molecules on the activity of PfFΔ40, fumarate was used as the substrate at a concentration of 3 mM. The molecules were tested at a concentration of 0.5 mM. For estimating K_i_ values for DL-MSA and meso-tartrate, the initial velocity was measured at varying concentrations of malate (46 μM to 12 mM) / fumarate (24 μM to 25 mM) with DL-MSA/ meso-tartrate fixed at different concentrations. The mode of inhibition was inferred from the type of intersection pattern of lines in the Lineweaver-Burk plot. The Ki value for DL-MSA was obtained from a global fit of the data by nonlinear regression analysis to a competitive model for enzyme inhibition using GraphPadPrism5.

### Generation and phenotyping of ΔfumACB strain of E. coli

In order to generate a fumarate hydratase null strain of *E. coli,* a *fumB* null strain (JW4083-1, *fumB748 (del)::kan)* (57), derived from the *E. coli* strainBW25113, was obtained from Coli Genetic Stock Centre ( CGSC), Yale University, New Haven, USA (57). To remove the kanamycin cassette flanked by FRT sites at the *fumB* gene locus and to subsequently knockout *fumA* and *fumC,* standard protocols were followed (58) and this is described in Supplementary Methods and S1 Fig. Knockout of the genes was validated by PCR using genomic DNA of the mutant strain as template and appropriate oligonucleotides (S1 Table). M9 minimal medium agar plates containing either fumarate or malate (0.4 %) as the sole carbon source and supplemented with trace elements were used to check the phenotype of the strains Δ*fumACB, ΔfumA, ΔfumB* and Δ*fumC.* An equal number of cells of each of these strains were spread on both malate and fumarate-containing M9 agar plates and the growth phenotype was scored at the end of 48 h of incubation at 37 °C under aerobic conditions.

### Complementation of FH deficiency in ΔfumACB strain with PfFH and growth inhibition with MSA

The Δ*fumACB* strain of *E. coli* was transformed with the plasmids pQE-PfFHFL, pQE-PfFΔ40, pQE-PfFΔ120, and pQE30 and selected on LB plate containing 100 μg ml^-1^ ampicillin and 50 μg ml^-1^ kanamycin. A single colony from the plate was inoculated into 10 ml LB broth and allowed to grow overnight. An aliquot of each of the cultures was washed three times with sterile M9 medium to remove traces of LB broth. The cells were resuspended in M9 medium and an OD600-normalized aliquot of the suspensions was spread on an M9 agar plate containing the appropriate carbon source and antibiotics. It should be noted that the parent strain BW25113 has a single copy of *lacI^+^* allele and not *lacI^q^* (59) and hence, for induction of protein expression in this strain using pQE30 based constructs, the addition of IPTG is optional.

To check the effect of MSA on the *E. coli strain ΔfumACB* with the plasmid pQE-PfFΔ40, the culture was grown overnight in 10 ml LB medium and 1 ml of the culture was washed twice with M9 minimal medium and resuspended in 1 ml M9 minimal medium. 150 μl of this suspension was added to tubes containing 5 ml of M9 minimal medium with 10 mM fumarate as the sole carbon source and appropriate antibiotics (50 μg ml^-1^kanamycin and 100 μg ml^-1^ampicillin). Varied concentrations of DL-MSA ranging from 1 μM to 15 mM were added to the tubes. The growth of the cultures was monitored by measuring OD_600_ at the end of 10 h.

### P. falciparum culture, transfection and growth inhibition with MX4

Intraerythrocytic stages of *P. falciparum* 3D7 strain (procured from MR4) were grown by the method established by Trager and Jensen (60). The parasites were grown in medium containing RPMI-1640 buffered with 25 mM HEPES and supplemented with 20 mM sodium bicarbonate, 0.5 % AlbuMAX I, 0.5 % glucose and 100 μM hypoxanthine. O positive erythrocytes from healthy volunteers were added to the culture to a final hematocrit of 2 % for regular maintenance. For examining the localization of PfFH, PM1KO strain (56) was transfected with the plasmid pGDB-PfFH. For this, preloading of erythrocytes (61) with plasmid DNA was carried out by electroporation using a square wave pulse (8 pulses of 365 V each lasting for 1 ms with a gap of 100 ms) program in BioRad-XL electroporator. Briefly, 100 μg of plasmid DNA dissolved in cytomix (61) was used for transfection of uninfected erythrocytes resuspended in cytomix. After electroporation, the cells were washed with incomplete media to remove cell debris and 1 ml of infected erythrocytes (2 % hematocrit and 6-8 % parasitemia) containing late schizont stage parasites was added. The parasites were allowed to reinvade and when the parasitemia reached 6-8 %, drug selection was started by the addition of trimethoprim (10 μM) and blasticidin S (2.5 μg ml^-1^). The strain of *P. falciparum* is referred to as PfFH-GFP. The strain was subjected to three rounds of drug cycling which included growing the cultures on and off blasticidin (10 days each) in the continuous presence of trimethoprim following which the genotyping of the strain was performed by PCR using genomic DNA as template and primers P1-P4 (S1 Table).

The IC_50_ value for DL-MSA for inhibition of parasite growth was determined by a serial twofold dilution. Briefly, the effect of DL-MSA on the viability of the 3D7 strain of *P. falciparum* was determined by counting the number of parasites in at least 1000 erythrocytes in Giemsa stained smears of cultures grown in the presence of increasing concentrations (3 μM-40 mM) of the drug.

### Mitochondrial staining and microscopy

For mitochondrial staining of PfFH-GFP parasites, the culture containing mixed stages of parasites was washed with incomplete medium twice to remove any traces of AlbuMAX I, the cells resuspended with incomplete medium containing 100 nM MitoTracker CM-H_2_XRos and incubated at 37 °C in a candle jar for 30 minutes. For nuclear staining, Hoechst 33342 was added to the culture to a final concentration of 5 μg ml^-1^ and incubated for an additional 5 minutes at 37 °C. For imaging, 500 μl of this culture was washed with incomplete medium once and the cells were resuspended in 50 % glycerol/PBS solution. 5 μl of the suspension was placed under a coverslip and imaged using DeltaVision Elite widefield microscope, GE, USA at room temperature. The images were processed using ImageJ (62, 63).

### P. berghei culturing and genetic manipulation

Intraerythrocytic asexual stages of *P. berghei* ANKA (procured from MR4) were maintained in BALB/c mice. For the generation of the knockout construct and for the transfection of parasites, established procedures were followed (64, 65). All transfection experiments were performed twice. Starting from *fh* genomic library clone (Clone ID: PbG01-2466a09) obtained from PlasmoGEM repository (Wellcome Trust Sanger Institute, UK), the *fh* gene knockout construct was generated by using the recombineering strategy described by Pfander et al. (32) (S2 Fig). This construct has, flanking the resistance marker, 1395 bp and 2049 bp DNA segments corresponding to regions upstream and downstream, respectively of the *fh* gene to enable gene knockout by double-crossover recombination. For transfection of *P. berghei,* the parasites were harvested from infected mice at a parasitemia of around 1-3 %. Around 0.8-1.0 ml of blood was obtained from each mouse and the parasites were synchronized at schizont stage by *in vitro* growth at 36.5 °C with constant shaking at an optimal speed of 120-150 rpm in medium containing RPMI-1640 with glutamine, 25 mM HEPES, 10 mM NaHCO_3_ and 20 % fetal bovine serum under a gassed environment (5 % oxygen, 5 % carbon dioxide and 90 % nitrogen). The schizonts were purified by density gradient centrifugation on Nycodenz and transfected with NotI digested linear DNA of the *fh* gene knockout construct using a 2D-nucleofector (Lonza, Switzerland). Pyrimethamine selection was started 1 day after transfection (65). Limiting dilution cloning of the drug resistant parasites was performed using 16 mice in two batches (32 mice in total). Genomic DNA was isolated from 17 individual parasite lines and subjected to series of diagnostic PCRs to check integration and loss of *fh* gene. Mouse strain dependent essentiality of *fh* for *P. berghei* was examined by transfection of wild-type parasites harvested from infected BALB/c mouse with *fh* gene knockout construct. An equal volume of parasite suspension from this single transfection reaction was injected intravenously into BALB/c and C57BL/6 mice. Genotyping of drug selected parasites obtained from both mice was performed by PCR using gene-and integration-specific oligonucleotides. The sequences of the oligonucleotides used are provided in S1 Table.

### Ethics statement

All animal experiments involving BALB/c and C57BL/6 mice adhered to the standard operating procedures prescribed by the Committee for the Purpose of Control and Supervision of Experiments on Animals (CPCSEA), a statutory body under the Prevention of Cruelty to Animals Act, 1960 and Breeding and Experimentation Rules 1998, Constitution of India. The study was a part of the project numbered HB004/201/CPCSEA and is approved by the Institutional animal ethics committee (IAEC) that comes under the purview of CPCSEA. Whole blood for *P. falciparum* culturing was collected from healthy volunteers with written informed consent.

## Acknowledgments

This project was funded by; 1) Department of Biotechnology, Ministry of Science and Technology, Government of India. Grant number: BT/PR11294/BRB/10/1291/2014 and BT/PR13760/COE/34/42/2015, 2) Science and Engineering Research Board, Department of Science and Technology, Government of India. Grant number: EMR/2014/001276 and, 3) Institutional funding from Jawaharlal Nehru Centre of Advanced Scientific Research, Department of Science and Technology, India VJ acknowledges CSIR for junior and senior research fellowships. AS acknowledges UGC for junior and senior research fellowships.

### Conflict of interest

The authors declare that they have no conflicts of interest with the contents of this article.

### Author contributions

VJ, AS, HB conceived and designed the experiments. VJ, AS, PK, JK performed the experiments. VJ, AS, HB analyzed the data. VJ, AS, HB wrote the paper:.

## FOOTNOTES

**Present addresses**: Current affiliation of JK-Molecular characterization/Analytical group, Biocon Research Ltd.-SEZ unit, Biocon Park, Bommasandra-Jigani link road, Bengaluru, 560099, INDIA.

